# Phylodynamic estimation of the within-host mutation rate of extended-spectrum beta-lactamase-producing Enterobacterales

**DOI:** 10.1101/2025.01.31.635889

**Authors:** Etthel M. Windels, Lisandra Aguilar-Bultet, Isabelle Vock, Laura Maurer Perkerman, Sarah Tschudin-Sutter, Tanja Stadler

## Abstract

Despite their clinical relevance, the within-host evolution of extended-spectrum beta-lactamase-producing Enterobacterales (ESBL-PE) is still poorly understood. To estimate the within-host mutation rates of ESBL-producing *Escherichia coli* and *Klebsiella pneumoniae* species complex, we fitted phylodynamic models to genomic sequence data of longitudinally collected rectal swabs from 64 colonized hospital patients. We estimated an average within-host mutation rate of 7.71e-07 [4.60e-07,1.10e-06] mutations/site/year for *E. coli* and 4.20e-07 [1.57e-07,7.14e-07] mutations/site/year for *K. pneumoniae* species complex, with limited variation observed across patients and sequence types. These estimates are presumably the most accurate reported so far and are useful for future epidemiological and evolutionary studies.

**Impact statement:** Understanding the within-patient mutation rates of extended-spectrum beta-lactamase (ESBL)-producing *E. coli* and *K. pneumoniae* species complex is critical for elucidating their evolutionary dynamics and transmission potential. In this study, we employed Bayesian phylodynamic models to analyze longitudinal isolates collected over a 10-year period from hospitalized patients. Our findings reveal limited variability in mutation rates across patients and sequence types, suggesting evolutionary stability within these pathogens in hospital settings. This work provides new insights into the persistence and evolutionary dynamics of ESBL-PE, offering valuable guidance for antimicrobial resistance surveillance and infection prevention strategies.

**Data summary:** The data used in this study are publicly available in the NCBI database under the BioProject number PRJNA910977. Supporting metadata are provided in Appendix 1. The code for the phylodynamic analyses, including the BEAST2 XML files, is available at https://github.com/EtthelWindels/esbl-pe_mutation. All supporting data, code and protocols have been provided within the article, supplementary data files or public repositories.

## Introduction

Extended-spectrum beta-lactamase-producing Enterobacterales (ESBL-PE) are clinically significant microorganisms associated with increased morbidity, mortality, and healthcare costs due to their resistance to a wide range of beta-lactam antibiotics, thereby complicating treatment options (1, 2). Understanding the transmission dynamics of these pathogens, particularly over long time periods, is critical to mitigate their impact. To investigate potential transmission events, it is essential to account for the diversity and evolution of pathogens within an individual host. In hospital settings, sequences from strains collected at different time points and body sites are valuable for such investigations. In a previous study (3), we demonstrated the long persistence of ESBL-PE strains within patients, with very low genomic variation over long time periods. We also provided preliminary estimates of the within-host mutation rates based on the number of single nucleotide polymorphisms (SNPs) identified between consecutive isolates relative to the first isolate, and the time span between the paired isolates. However, this method did not consider the phylogenetic tree, thereby potentially underestimating the true divergence times and hence overestimating the mutation rates. In this study, we aimed to overcome this limitation by using Bayesian phylodynamic models, which consider the phylogenetic tree as well as an evolutionary model accounting for unobserved mutations (including back mutations, multiple mutations at the same site, and parallel mutations), to obtain more accurate estimates of the within-patient mutation rates of ESBL-producing *E. coli* and *K. pneumoniae* species complex collected during a 10-year longitudinal study.

## Results

### Estimation of average within-patient mutation rates

In a previous study at the University Hospital Basel, Switzerland, 189 rectal swab isolates were longitudinally collected from 64 patients colonized with ESBL-PE (3). We used these data to generate per-patient sequence alignments and fitted a constant population coalescent model with strict clock, assuming a shared within-patient mutation rate for all patients per species. Only patients with at least three serial isolates belonging to the same strain (28/64 patients and 107/189 isolates) were included in the analysis. Inference was done in a Bayesian setting, meaning that we inferred posterior distributions which reflect the information in the data about each parameter, in combination with the prior distribution. The posterior mean for the average mutation rate was estimated to be 6.04e-07 (95% highest posterior density interval, HPDI: [3.69e-07,8.40e-07]) mutations/site/year for *E. coli* and 3.75e-07 (95% HPDI: [1.77e-07,5.83e-07]) mutations/site/year for *K. pneumoniae* species complex (Figure 1). These estimates are slightly lower than the estimates reported before (1.4e-06 (interquartile range, IQR: [7.6e-07,3.2e-06]) mutations/site/year for *E. coli* and 1.5e-06 (IQR: [7.3e-07,3.5e-06]) mutations/site/year for *K. pneumoniae* species complex) (3). When all patients – including those with only two serial isolates – were considered, the posterior estimates increased slightly (8.69e-07 [6.21e-07,1.09e-06] mutations/site/year for *E. coli* and 4.70e-07 [2.41e-07,7.14e-07] mutations/site/year for *K. pneumoniae* species complex) and the posterior distributions more closely followed the prior (Figure S1), suggesting that the isolates of these patients are uninformative about the mutation rate and may even slightly bias the posterior estimates in combination with the chosen prior.

**Figure 1:**
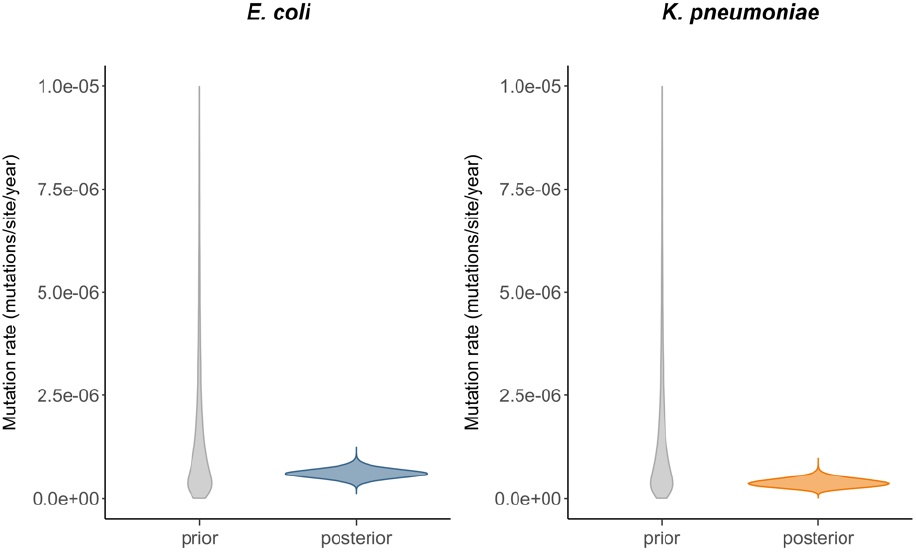
Average within-patient mutation rate estimates. Prior (grey) and posterior (colored) distributions of the within-patient mutation rate, averaged over all patients for which at least three serial isolates were available (21 patients with 78 isolates for *E. coli* and 7 patients with 29 isolates for *K. pneumoniae* species complex). The posterior mean and 95% HPDI correspond to 6.04e-07 [3.69e-07,8.40e-07] mutations/site/year for *E. coli* and 3.75e-07 [1.77e-07,5.83e-07] mutations/site/year for *K. pneumoniae* species complex.

### Estimation of patient-specific within-patient mutation rates

We then adjusted the phylodynamic model to infer a within-host mutation rate per patient. In addition to the average mutation rate informed by all patient alignments, this model was further parametrized with patient-specific mutation rate multipliers, with each patient-specific mutation rate corresponding to the product of the average mutation rate and the patient-specific multiplier. The phylogenetic trees inferred under this model are shown in Figure S2. The resulting estimates for the average mutation rate were similar to the estimates from the previous analysis (7.71e-07 (95% HPDI: [4.60e-07,1.10e-06]) mutations/site/year for *E. coli* and 4.20e-07 (95% HPDI: [1.57e-07,7.14e-07]) mutations/site/year for *K. pneumoniae* species complex). The posterior estimates for the patient-specific multipliers were close to one for most patients, suggesting limited patient-to-patient variability in within-host mutation rates (Figure 2). When additionally considering patients for which only two isolates were available, the estimates for the average mutation rate again increased slightly. Consequently, the patient-specific mutation rate for some patients was estimated to be lower than this average (Figure S3).

**Figure 2:**
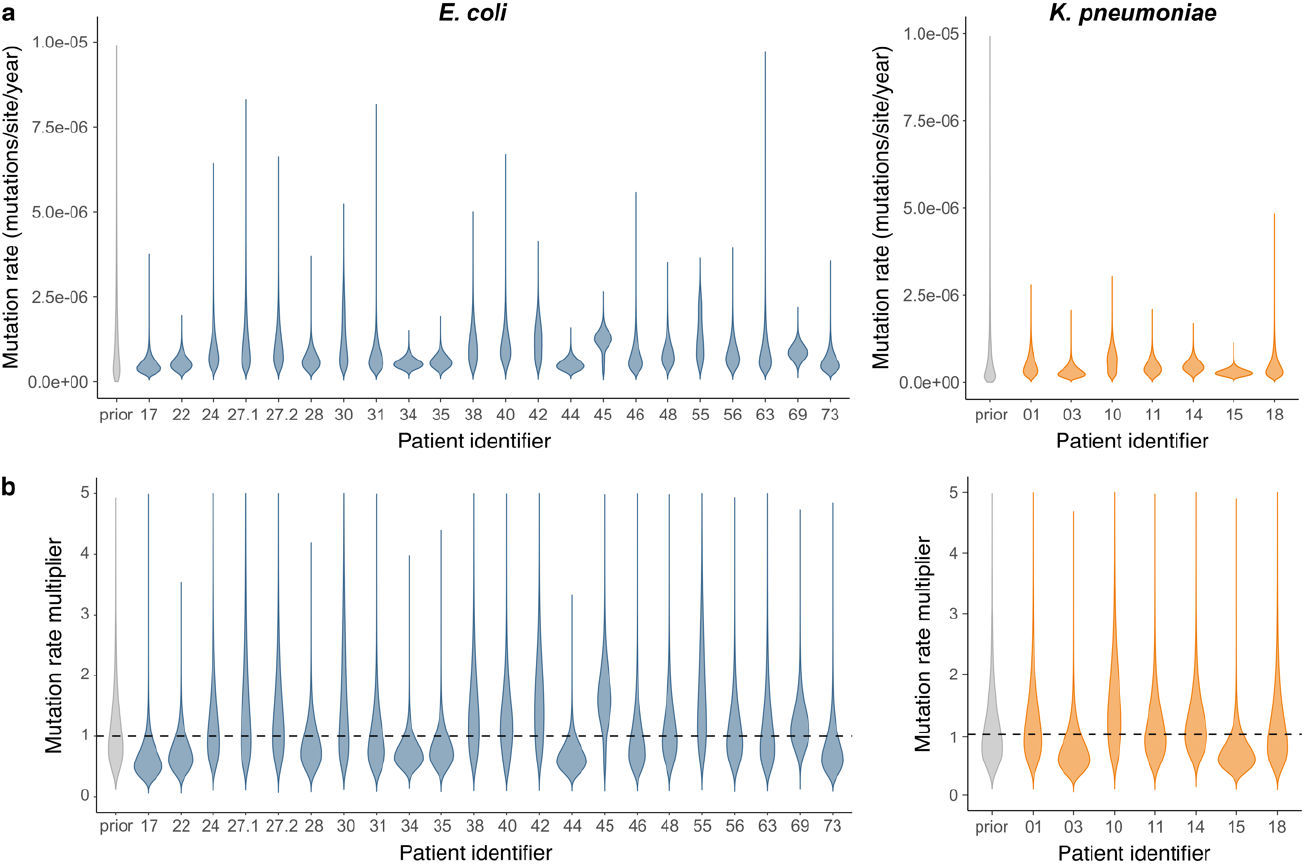
Patient-specific within-patient mutation rate estimates. a) Prior (grey) and posterior (colored) distributions of patient-specific within-patient mutation rates, estimated for all patients for which at least three serial isolates were available (21 patients with 78 isolates for *E. coli* and 7 patients with 29 isolates for *K. pneumoniae* species complex). Each patient-specific mutation rate estimate corresponds to the product of the average mutation rate estimate (7.71e-07 [4.60e-07,1.10e-06] mutations/site/year for *E. coli* and 4.20e-07 [1.57e-07,7.14e-07] mutations/site/year for *K. pneumoniae* species complex) and a patient-specific multiplier estimate. b) Prior (grey) and posterior (colored) distributions of patient-specific mutation rate multipliers. The posterior estimates are close to one for most patients, suggesting limited patient-to-patient variability. Patient identifiers 27.1 and 27.2 correspond to the same patient but different strains, so two mutation rates were estimated for this patient.

**Figure 3:**
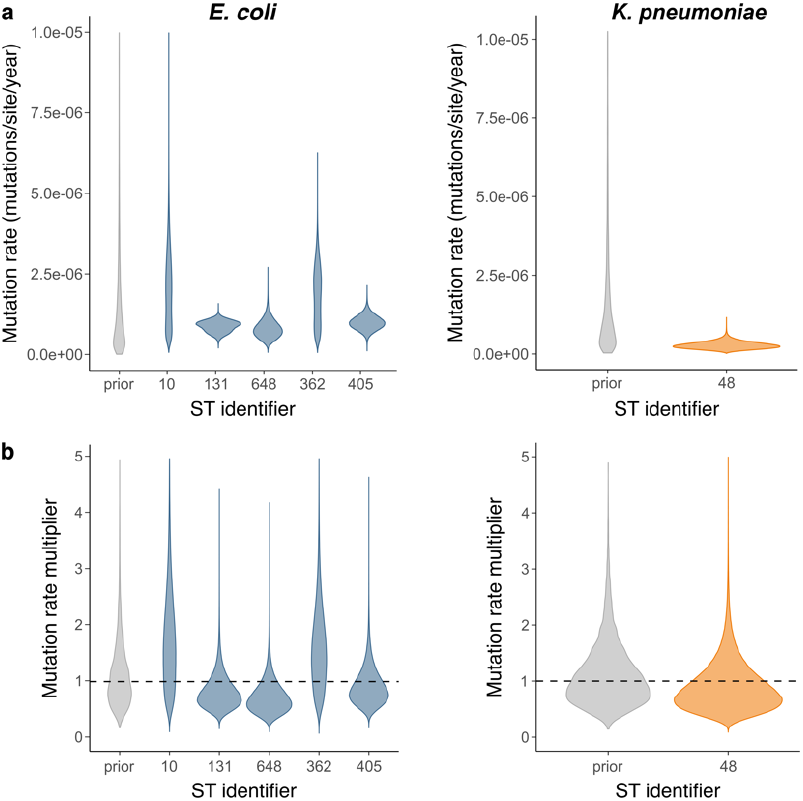
ST-specific within-patient mutation rate estimates. a) Prior (grey) and posterior (colored) distributions of ST-specific within-patient mutation rates, estimated for all STs with at least two patients, of which at least one patient with at least three serial isolates (30 patients with 87 isolates belonging to 5 STs for *E. coli* and 3 patients with 10 isolates belonging to 1 ST for *K. pneumoniae* species complex; Table S2). Each ST-specific mutation rate estimate corresponds to the product of the average mutation rate estimate (1.17e-06 [4.73e-07,1.94e-06] mutations/site/year for *E. coli* and 3.40e-07 [3.97e-08,7.65e-07] mutations/site/year for *K. pneumoniae* species complex) and a ST-specific multiplier estimate. b) Prior (grey) and posterior (colored) distributions of ST-specific mutation rate multipliers. The posterior estimates are close to one for most STs, implying that the data do not support an association between ST and mutation rate.

We tested the robustness of our estimates to prior assumptions, by setting a different prior distribution on the average mutation rate (Figure S4) and by assuming an exponential growth coalescent model (Figure S5). In both cases, the mutation rate estimates were almost unaffected. When patient-specific mutation rates were estimated independently instead of through a shared mutation rate combined with independent multipliers, the overall trends remained the same, although with increased uncertainty in the posterior estimates (Figure S6).

### Estimation of sequence type-specific within-patient mutation rates

To check for an association between sequence types (STs) and within-patient mutation rates, we inferred a mutation rate for each ST with at least two sampled patients, of which at least one patient had at least three serial isolates (Table S2). Similar to the patient-specific mutation rate inference, we generated per-ST alignments and inferred an average mutation rate informed by all ST alignments as well as ST-specific mutation rate multipliers. The average mutation rate estimate for *E. coli* (1.17e-06 (95% HPDI: [4.73e-07,1.94e-06]) mutations/site/year) was slightly higher than in previous analyses while the estimate for *K. pneumoniae* species complex (3.40e-07 (95% HPDI: [3.97e-08,7.65e-07]) mutations/site/year) was similar. However, the patient set used in this analysis is slightly different from the one used in previous analyses and the number of patients per ST varies, making it difficult to predict the effect of clustering by ST on the average mutation rate estimate. For *K. pneumoniae* species complex, only one ST fulfilled the inclusion criteria, making the multiplier estimate redundant. For *E. coli*, the ST-specific mutation rate multiplier estimates did not deviate from one and showed limited variability, implying that the data do not support an association between the within-patient mutation rate and ST. Including all patients in the analysis resulted in a slight increase in the estimated average mutation rate for *K. pneumoniae* species complex, and again limited ST-to-ST variability (Figure S7).

## Discussion

In this study, we used phylodynamic models to estimate within-patient mutation rates of ESBL-producing *E. coli* and *K. pneumoniae* species complex from longitudinal isolates collected in a hospital setting. These mutation rates were estimated as an average over all patients as well as on a per-patient and per-ST basis, using models that were parametrized such that the information in the available data was used as efficiently as possible. The resulting estimates of the average within-patient mutation rate were slightly lower than reported previously (3). These previous estimates were obtained by dividing the number of observed SNPs by the time-based sampling interval, for each isolate with respect to the first isolate of the patient, and taking the average over all pairs. By considering only the sampling interval instead of the total time since the most recent common ancestor of all isolates per patient, the evolutionary time is likely underestimated, which might explain the higher mutation rate estimates obtained in the previous study (3). Another difference is that our phylodynamic models account for unobserved mutations such as back mutations. If many such unobserved mutations occurred, ignoring them could result in an overestimation of the evolutionary rate. For *E. coli*, the new estimates derived in this study correspond well to the previously reported within-host mutation rate of the non-pathogenic *E. coli* ED1a clone (6.90e-7 mutations/site/year) estimated from a single healthy individual using a similar Bayesian method (4).

The limited variation observed across patients was not entirely surprising, given the similar backgrounds of the majority of study participants in terms of medical history and previous comorbidities. The limited across-ST variation suggests that differences in adaptive potential are not directly caused by differences in mutation rate, not even for the globally spread ST131 of *E. coli*.

The per-patient mutation rate was estimated in two different ways, either by estimating the mutation rate independently for each patient alignment (with the same, relatively broad prior distribution set on each individual mutation rate), or by combining all alignments to inform a shared mutation rate (on which a broad prior was set) and including patient-specific multipliers (with a narrow prior around one) in the model to allow for patient-to-patient variation. The second method resulted in lower posterior uncertainty, likely because it uses the available data more efficiently by combining information from multiple patients while allowing for patient differences.

A limitation of this study was that only a few isolates were available for most patients, resulting in rather large posterior uncertainty in patient-specific estimates. Our analyses showed that patients with less than three isolates are better excluded, as they result in more noisy and potentially skewed estimates given our prior choices. Future studies including more patients, and particularly more isolates per patient, have the potential to further refine the within-host mutation rates reported here. Furthermore, including environmental samples from longitudinal studies would help bridge the gap in the evolutionary history of these microorganisms within a One Health context.

## Materials and Methods

### Study population

We used previously sequenced serial isolates from an observational cohort study at the University Hospital Basel, Switzerland (3). Patients included in this study were admitted to the hospital between January 2008 and December 2018 and had ESBL-PE belonging to the same species (*E. coli* or *K. pneumoniae* species complex) detected in at least two consecutive rectal swabs. In total, 189 rectal swab isolates were collected, comprising 134 *E. coli* isolates from 47 patients and 55 *K. pneumoniae* species complex isolates from 19 patients. Two patients (patient 17 and 18) harbored both species. The isolates included in this study along with their associated patients, ST classification and cluster assignments (i.e., isolates within the same cluster belong to the same strain) are summarized in Appendix 1.

### Multiple sequence alignments

Raw Illumina sequencing reads were processed as described before (3). Per-patient alignments were generated using Snippy v.4.6.0 (https://github.com/tseemann/snippy). A patient-specific reference was generated by concatenating all contigs from the first sample of each patient. Two *E. coli* patients harbored two different strains, therefore two different alignments and references were generated for each patient. The query samples were taken from the consensus of each pairwise mapping and were concatenated in the same way. From the complete alignments (4,714,797 sites for *E. coli*; 5,654,681 sites for *K. pneumoniae* species complex), only the variable sites were kept. Only patients for which at least three serial isolates belonging to the same strain were available were retained for the main analyses. STs were identified using Ridom SeqSphere+ v.6.0 (Ridom, Münster, Germany), and only STs containing at least one patient with at least three isolates were retained for the main analyses.

### Phylodynamic model

A constant population size coalescent model was fitted onto the set of per-patient sequence alignments, with patient-specific parameters for the effective population size. We further assumed a strict molecular clock and a general time-reversible nucleotide substitution model with four gamma rate categories to account for site-to-site rate heterogeneity (GTR+Γ_4_). All substitution model parameters were assumed to be shared across all patients. In a first analysis, the mutation rate was also assumed to be shared across all patients. In the follow-up analyses, the model was additionally parametrized with either patient-specific or ST-specific mutation rate multipliers. The overall patient-specific or ST-specific mutation rate was then equal to the shared mutation rate multiplied by the respective multiplier. All parameters and their prior distributions are listed in Table S1.

### Phylodynamic inference

We performed phylodynamic inference using the feast package v8.3.1 (https://github.com/tgvaughan/feast/releases/tag/v8.3.1) in BEAST v2.7.6 (5, 6). Data from each species (*E. coli* and *K. pneumoniae* species complex) were analyzed independently. Variable SNP alignments were augmented with a count of invariant A, C, G, and T nucleotides (7). For each analysis, three independent Markov Chain Monte Carlo chains were run, with states and trees sampled every 100,000 steps. Convergence was assessed with Tracer (8), confirming that the effective sample size (ESS) was at least 200 for the parameters of interest. 10% of each chain was discarded as burn-in, and the remaining samples across the three chains were pooled with LogCombiner (6), resulting in at least 50,000,000,000 iterations in combined chains.

### Sensitivity analyses

The robustness of the phylodynamic inference to prior assumptions was tested by setting a Lognormal(-14.60,1.25) prior (corresponding to a mean in real space of 10e-6) on the mutation rate, as well as by using an exponential growth coalescent model with patient-specific growth rate parameters and a Laplace(0.001,0.5) prior distribution on these growth rates. Moreover, we tested the effect of including patients for which only two isolates are available. Finally, instead of estimating a shared mutation rate and patient-specific multipliers, we tested the performance of a model with independent patient-specific mutation rate parameters, with the same Lognormal(-13.82,1.25) prior distribution set on each mutation rate.

## Author contributions

Conceptualization: S.T.S., L.A.-B., T.S., E.M.W.; Data curation: L.A.-B., I.V, L.M.P.; Formal analysis: E.M.W., L.A.-B.; Funding acquisition: S.T.S., T.S.; Investigation: E.M.W., L.A.-B., T.S., S.T.S.; Methodology: E.M.W., T.S.; Supervision: T.S., S.T.S.; Visualization: E.M.W.; Writing – original draft: E.M.W., L.A.-B.; Writing – review and editing: E.M.W., L.A.-B., I.V, L.M.P., T.S., S.T.S.

## Acknowledgements

Calculations were performed on the Euler cluster at ETH Zürich and sciCORE (http://scicore.unibas.ch/) scientific computing center at the University of Basel. We would like to thank Timothy G. Vaughan for help with the phylodynamic analyses and Louis du Plessis for insightful discussions.

