## Supplementary Material for "Phylodynamic estimation of the within-host mutation rate of extended-spectrum beta-lactamase-producing Enterobacterales"

### Supplementary Tables

**Table S1:** Prior distributions for the parameters of the phylodynamic model (see Materials and Methods for more information about the individual parameters)

| Parameter | Prior |
|---|---|
| Effective population size | 1/X |
| Shared mutation rate | Lognormal(-13.82,1.25) |
| Mutation rate multiplier | Lognormal(0,0.5) |
| Gamma shape | Exp(1) |
| AC substitution rate | Gamma(0.05,10) |
| AG substitution rate | Gamma(0.05,20) |
| AT substitution rate | Gamma(0.05,10) |
| CG substitution rate | Gamma(0.05,10) |
| GT substitution rate | Gamma(0.05,10) |
| Proportion of invariant sites | Uniform(0,1) |

**Table S2:** Sequence types (STs) and corresponding patients used in the main analyses. Only STs with at least two patients, of which at least one patient had at least three serial isolates, were included. Patient identifiers 30.1 and 30.2 correspond to the same patient but different strains.

| **Species** | **ST** | **Patient identifiers** |
|---|---|---|
| *E. coli* | 10 | 18, 61, 63 |
| *E. coli* | 131 | 21, 24, 26, 27, 28, 29, 32, 34, 35, 38, 39, 40, 41, 43, 44, 45, 62, 66, 71 |
| *E. coli* | 362 | 30.1, 30.2, 60 |
| *E. coli* | 405 | 46, 52, 69 |
| *E. coli* | 648 | 22, 51, 55 |
| *K. pneumoniae* species complex | 8 | 08, 15, 17 |

### Supplementary Figures

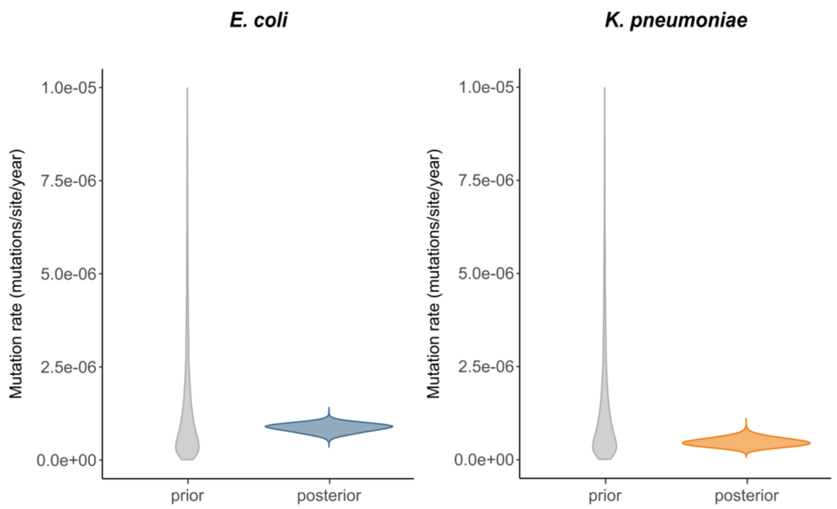


**Figure S1: Average within-patient mutation rate estimates on all patients.** Prior and posterior distributions of the within-patient mutation rate, averaged over all patients (including those with only two serial isolates). The posterior mean and 95% HPDI correspond to 8.69e-07 [6.21e-07,1.09e-06] mutations/site/year for *E. coli* and 4.70e-07 [2.41e-07,7.14e-07] mutations/site/year for *K. pneumoniae* species complex.

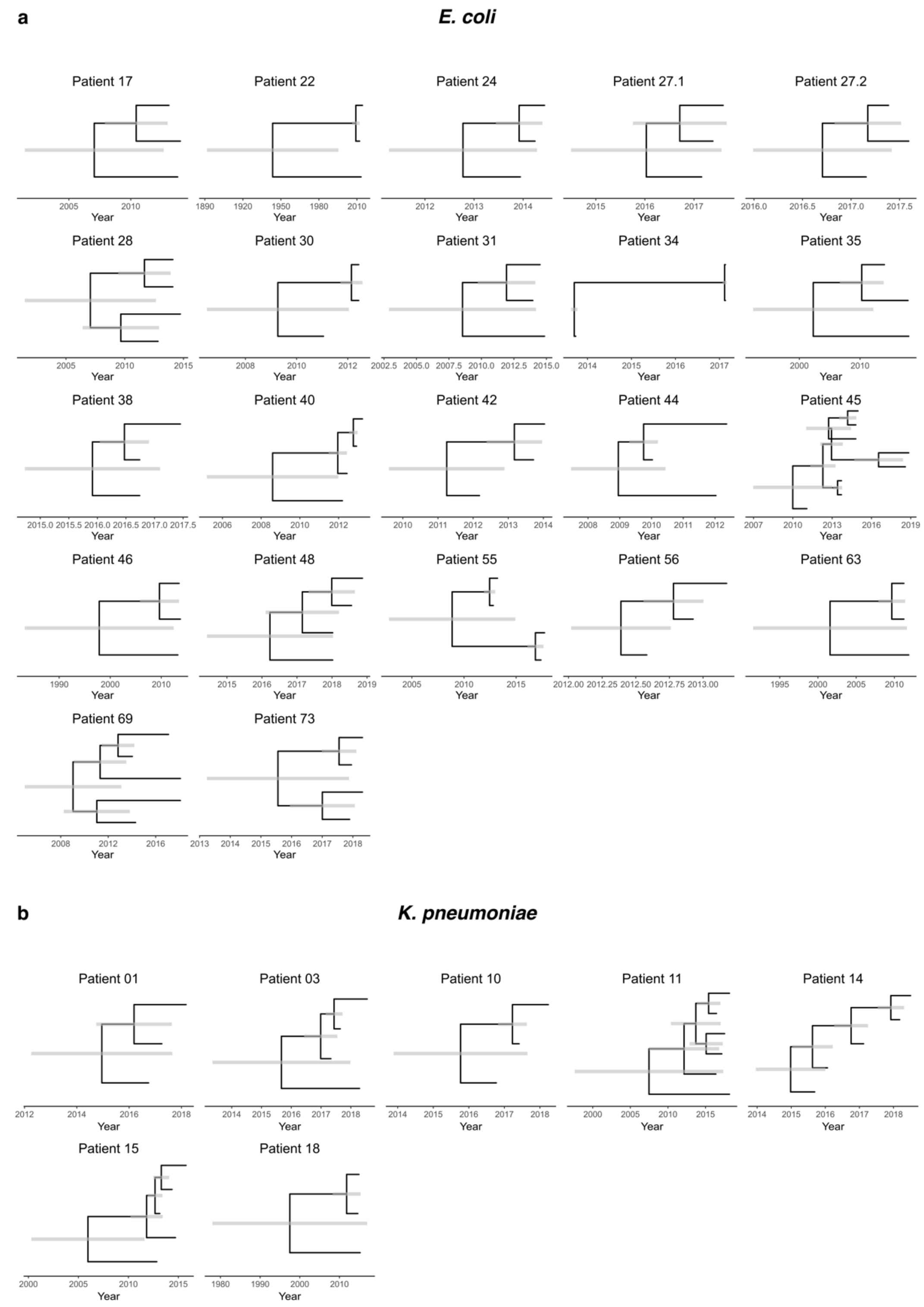


**Figure S2: Posterior maximum clade credibility trees.** Posterior maximum clade credibility trees per species and per patient, summarizing the posterior tree distribution resulting from the phylodynamic analyses on all patients for which at least three *E. coli* (a) or *K. pneumoniae* species complex (b) isolates were available. Grey shades represent 95% HPDIs on the node heights. Patient identifiers 27.1 and 27.2 correspond to the same patient but different strains.

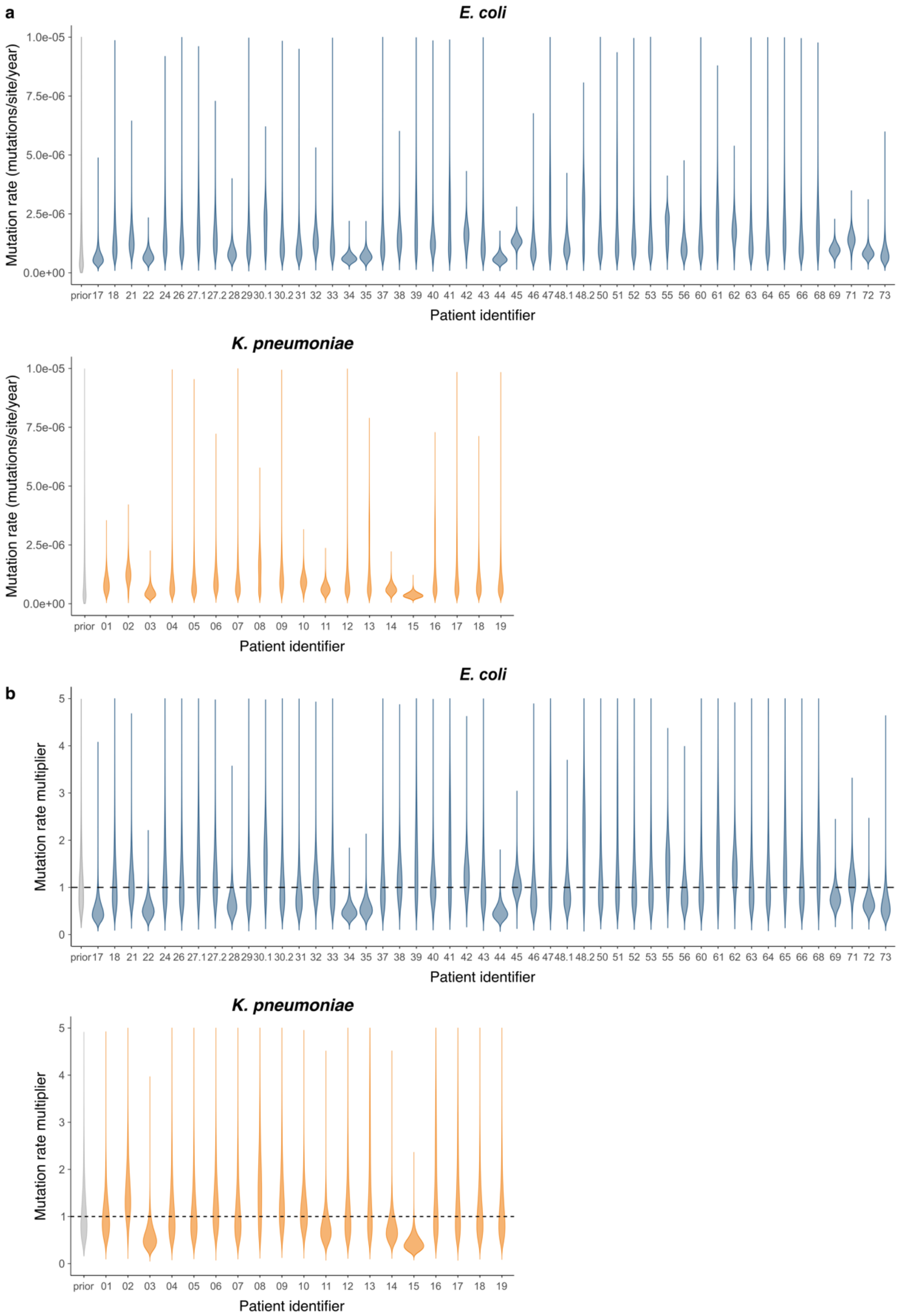


**Figure S3: Patient-specific within-patient mutation rate estimates on all patients.** a) Prior (grey) and posterior (colored) distributions of patient-specific within-patient mutation rates, estimated for all patients (including those with only two serial isolates). Each patient-specific mutation rate estimate corresponds to the product of the average mutation rate estimate (1.28e-06 [8.72e-07,1.69e-06]

mutations/site/year for *E. coli* and 8.69e-07 [3.85e-07,1.40e-06] mutations/site/year for *K. pneumoniae* species complex) and a patient-specific multiplier estimate. b) Prior (grey) and posterior (colored) distributions of patient-specific mutation rate multipliers. Patient identifiers 27.1/27.2, 30.1/30.2, and 48.1/48.2 correspond to the same patient but different strains, so two mutation rates were estimated for these patients. 30.1 and 48.1 correspond to the strains included in the main analyses. Patients 17 and 18 harbored both *E. coli* and *K. pneumoniae*.

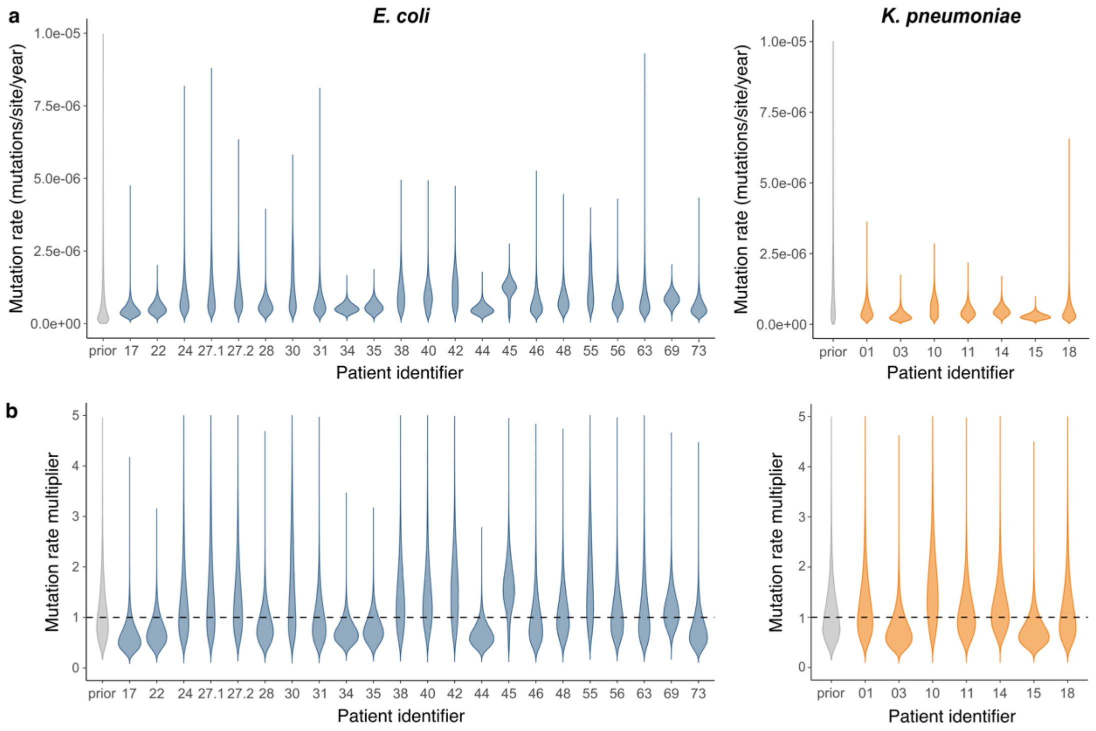


**Figure S4: Patient-specific within-patient mutation rate estimates, using a different mutation rate prior.** a) Prior (grey) and posterior (colored) distributions of patient-specific within-patient mutation rates, estimated for all patients for which at least three serial isolates were available, assuming a different prior distribution on the mutation rate (see Materials and Methods) as a sensitivity check. Each patient-specific mutation rate estimate corresponds to the product of the average mutation rate estimate (7.38e-07 [4.29e-07,1.07e-06] mutations/site/year for *E. coli* and 4.07e-07 [1.52e-07,6.95e-07] mutations/site/year for *K. pneumoniae* species complex) and a patient-specific multiplier estimate. b) Prior (grey) and posterior (colored) distributions of patient-specific mutation rate multipliers. All posterior distributions are close to those inferred under the main model (Figure 2), suggesting robustness to the choice of prior. Patient identifiers 27.1 and 27.2 correspond to the same patient but different strains, so two mutation rates were estimated for this patient.

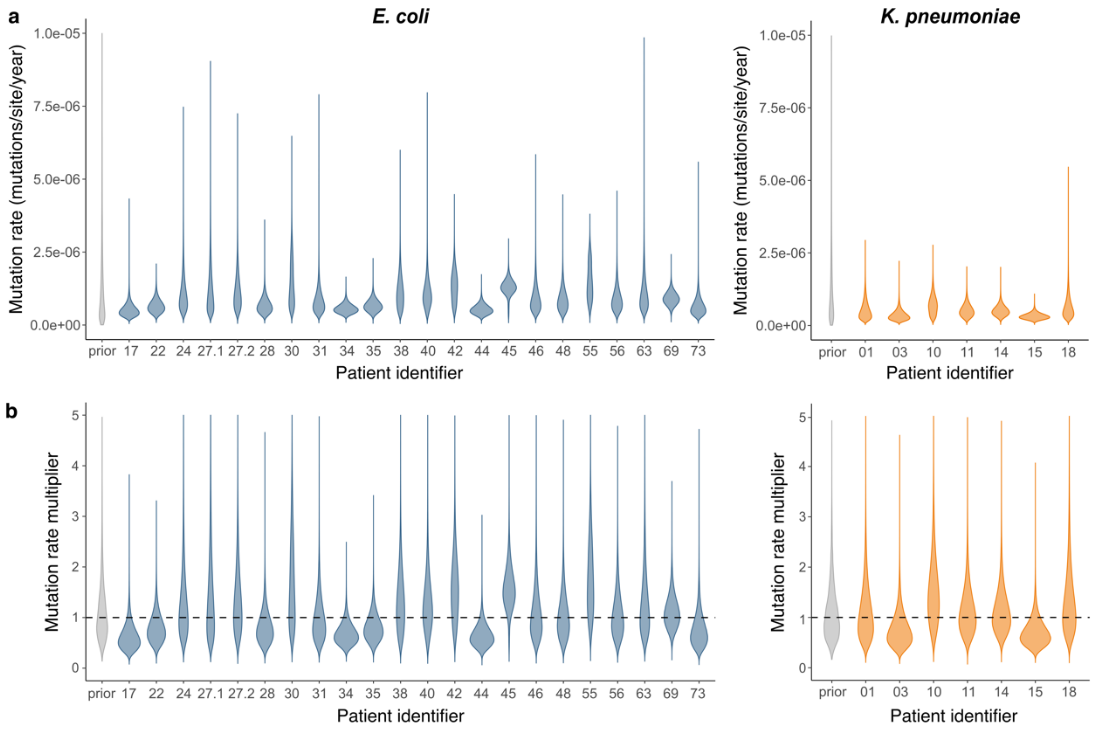


**Figure S5: Patient-specific within-patient mutation rate estimates, using an exponential growth coalescent model.** a) Prior (grey) and posterior (colored) distributions of patient-specific within-patient mutation rates, estimated for all patients for which at least three serial isolates were available, using an exponential growth coalescent model as a sensitivity check. Each patient-specific mutation rate estimate corresponds to the product of the average mutation rate estimate (8.39e-07 [5.28e-07,1.16e-06] mutations/site/year for *E. coli* and 4.76e-07 [2.05e-07,7.81e-07] mutations/site/year for *K. pneumoniae* species complex) and a patient-specific multiplier estimate. b) Prior (grey) and posterior (colored) distributions of patient-specific mutation rate multipliers. All posterior distributions are close to those inferred under the main model (Figure 2), suggesting robustness to the coalescent model. Patient identifiers 27.1 and 27.2 correspond to the same patient but different strains, so two mutation rates were estimated for this patient.

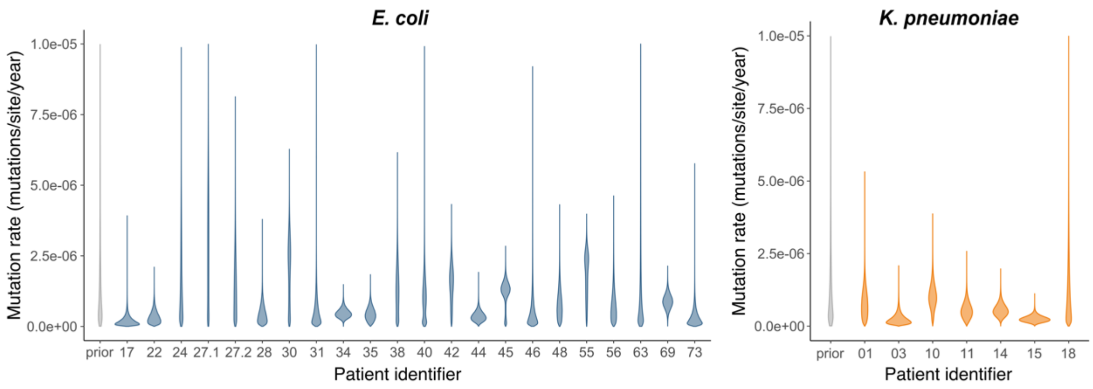


**Figure S6: Independent patient-specific within-patient mutation rate estimates.** Prior (grey) and posterior (colored) distributions of patient-specific within-patient mutation rates, estimated for all patients for which at least three serial isolates were available. In contrast to the main analysis, mutation rates were estimated independently for each patient. Patient identifiers 27.1 and 27.2 correspond to the same patient but different strains, so two mutation rates were estimated for this patient.

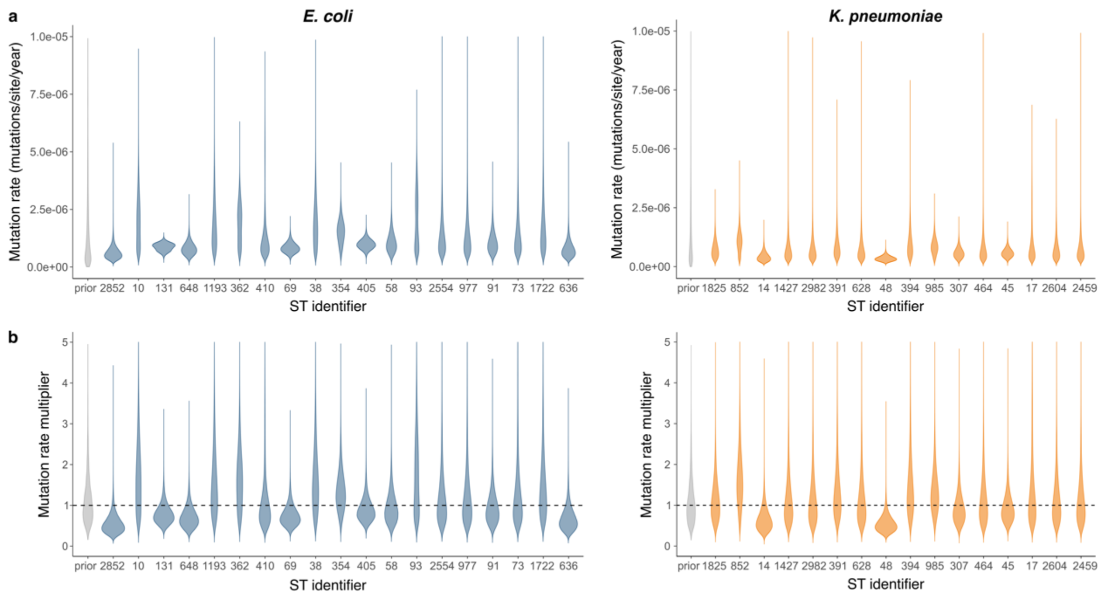


**Figure S7: ST-specific within-patient mutation rate estimates.** a) Prior (grey) and posterior (colored) distributions of ST-specific within-patient mutation rates, estimated for all STs on all available isolates. Each ST-specific mutation rate estimate corresponds to the product of the average mutation rate estimate (1.16e-06 [5.96e-07,1.76e-06] mutations/site/year for *E. coli* and 6.60e-07 [2.99e-07,1.05e-06] mutations/site/year for *K. pneumoniae* species complex) and a ST-specific multiplier estimate. b) Prior (grey) and posterior (colored) distributions of ST-specific mutation rate multipliers. The posterior estimates are close to one for most STs, implying that the data do not support an association between ST and mutation rate.
